# Coalescent-Based Time-Stratified Statistics Reveal Population Structure Dynamics using the Ancestral Recombination Graph

**DOI:** 10.64898/2026.08.11.744210

**Authors:** Yun Deng, Jonathan K. Pritchard, Jeffrey P. Spence

**Affiliations:** Department of Genetics, Stanford University; Department of Biology, Stanford University; Institute for Human Genetics, University of California, San Francisco; Department of Epidemiology & Biostatistics, University of California, San Francisco

## Abstract

Many questions in population genetics require reconstructing evolutionary history through time, such as inferring how population structure has changed throughout the past. Yet, many existing approaches have only an implicit temporal component, using quantities such as allele frequency or haplotype length as rough proxies for age. Recent advances in the inference of Ancestral Recombination Graphs (ARGs) have made it possible to estimate the entire sequence of local genealogies along the genome. These genealogies explicitly encode how samples are related to each other at different time points in the past, enabling the inference of how population structure has changed over time. To this end, recent work has used ARGs to define time-stratified versions of widely-used population genetics summary statistics in an attempt to capture the population structure present within a particular time window. Here, we show that naive approaches result in statistics that cannot isolate the population structure present solely within the targeted time window. To address this problem, we introduce a framework of **coalescent-based time-stratified statistics**, which use coalescence probabilities to partition classical summary statistics into interval-specific contributions. Using coalescent simulations, we demonstrate that these statistics accurately isolate population structure at different temporal depths and avoid spurious signals. Our results highlight the necessity of integrating coalescent theory into ARG-based temporal analyses and provide a principled and practical foundation for studying the dynamics of population structure through time.

## 1 Introduction

Many fundamental questions in population genetics are inherently temporal in nature. For example, demographic inference aims to reconstruct changes in population size, structure, and connectivity over time [Gutenkunst et al., 2009, Li and Durbin, 2011, Pickrell and Pritchard, 2012, Liu and Fu, 2015, Kamm et al., 2017]. Similarly, studies of natural selection seek to determine when selective pressures acted and how their strength changed over time [Smith et al., 2018, Stern et al., 2019, Vaughn and Nielsen, 2024, Irving-Pease et al., 2024]. Consequently, a wide range of methods can be viewed as implicitly performing **time-stratified analyses**, trying to understand the evolutionary forces acting at a particular moment in time.

Existing approaches, however, typically only indirectly stratify by time. For example, identity-by-descent (IBD) methods focus on very recent history by identifying long, shared haplotypes, which are unlikely to persist beyond a few generations [Browning and Browning, 2015, Fournier et al., 2023]. Other approaches, such as iHS, SDS and related statistics [Voight et al., 2006, Field et al., 2016], stratify variants by allele frequency as a heuristic proxy for allele age. While effective in specific regimes, these methods do not provide a principled, generalizable way to analyze evolutionary processes across arbitrary time depths, and their interpretation often depends on model-specific assumptions.

Meanwhile, there has been tremendous progress in the development of methods to infer Ancestral Recombination Graphs (ARGs). The ARG [Griffiths, 1981, Hudson, 1983] is a data structure that describes the full sequence of local genealogies along the genome for a sample of individuals. The ARG is a sufficient statistic for many population genetic parameters, containing all of the information that can possibly be extracted from a sample of individuals [Griffiths and Marjoram, 1997]. Despite its central role in coalescent theory, inferring the ARG from genomic data has long been challenging [Wiuf and Hein, 1999, McVean and Cardin, 2005]. Only in recent years have advances in statistical modeling, algorithms, and computational efficiency made realistic ARG inference feasible at scale [Rasmussen et al., 2014, Speidel et al., 2019, Kelleher et al., 2019, Wohns et al., 2022, Deng et al., 2025a,b].

The ARG naturally contains an explicitly temporal dimension. Each marginal tree encodes ancestral relationships at a specific locus, and branch lengths correspond directly to time [Hudson, 2002]. As a result, the ARG offers a time-resolved view of evolutionary history, in which population structure [Kelleher et al., 2019], demography [Speidel et al., 2019, Deng et al., 2025b], and selection [Vaughn and Nielsen, 2024] can in principle be examined at any chosen time depth. This perspective has motivated growing interest in extracting information from inferred ARGs in a time-stratified manner, with the goal of understanding how evolutionary processes vary through time [Nielsen et al., 2025].

A natural starting point for performing time-stratified analyses is to generalize classical population genetic summary statistics—such as the Genetic Relatedness Matrix (GRM) [Yang et al., 2011], *F_st_*[Wright, 1949, Weir and Cockerham, 1984], and *F*-statistics [Patterson et al., 2012]—to specific time intervals using the ARG. This approach promises to describe different aspects of the evolutionary process through time, and there have been recent efforts in this direction [Fan et al., 2022, Speidel et al., 2025].

Many classical summary statistics can be defined as functions of the genotype matrix, *G*, where *G_ik_* denotes the allelic state at site *k* in haplotype (or individual) *i*. For example, the GRM, *K*, for haplotype data is commonly defined as:

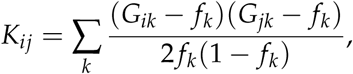

where *f_k_* is the derived allele frequency at site *k*. A centered, but unscaled version of the GRM is sometimes used instead,

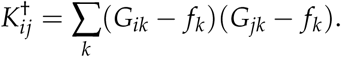

A naive approach to defining statistics over a specific time interval, say [*t_l_*, *t_u_*], is to restrict the genotype matrix to polymorphic sites whose underlying mutations arose within that time interval (Figure 1). Restricting to such sites results in a filtered genotype matrix, *G*^′^, on which standard summary statistics can be computed using their usual definitions. We refer to this approach as constructing **site-based time-stratified statistics**, consistent with the terminology used in Ralph et al. [2020].

**Figure 1:**
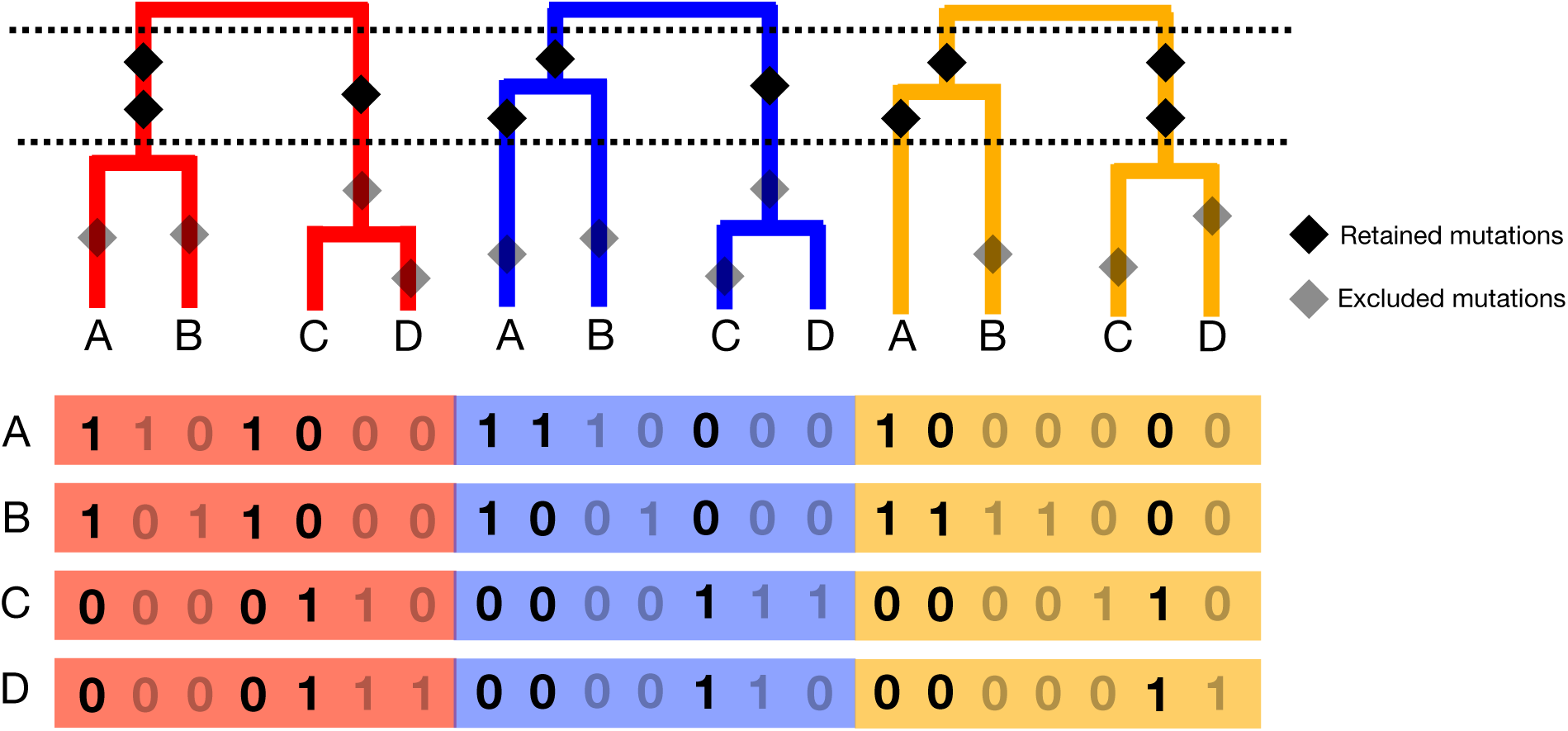
Site-based time-stratified statistics from the ARG. One approach to constructing a summary statistic for a given time interval (dashed lines) is to retain only polymorphic sites arising from mutations whose ages fall within the interval (dark diamonds), and to compute the statistic using the filtered genotype matrix with the standard definition.

This strategy closely mirrors the approach taken by the eGRM method [Fan et al., 2022], which defines time-stratified GRMs by focusing on mutations arising within specified time windows. eGRM further leverages the ARG by computing the *expected* GRM under a model in which mutations are placed uniformly along branches within the interval, rather than relying on observed mutations. This expectation-based formulation often leads to simplified expressions in terms of branch lengths and marginal tree topologies, enabling efficient computation via the tskit framework and in principle removes the noise induced by the randomness of the mutational process [Kelleher et al., 2016, Ralph et al., 2020, Wong et al., 2024, Jeffery et al., 2026].

We refer to this expectation-based approach—where statistics are computed by effectively “dropping” an infinite number of mutations onto ARG branches within a given time-interval—as **branch-based time-stratified statistics**. This approach was described in Ralph et al. [2020], and our naming of these statistics is consistent with their terminology.

Several recent methods build on this idea. For example, branch-PCA [Lehmann et al., 2025] performs principal component analysis using a branch-based, time-stratified (unscaled) GRM as a covariance matrix, implemented as TreeSequence.pca() in tskit. Similarly, Twigstats [Speidel et al., 2025] computes branch-based *F*-statistics over time intervals of the form [0, *t*], for arbitrary *t*, to characterize demographic history in the recent past.

### 1.1 Our contribution

Despite their intuitive appeal, here we show that both site-based and branch-based definitions of time-stratified statistics can be difficult to interpret and do not always isolate information from the intended time interval. In particular, even when mutations are strictly filtered by age or expectations are restricted to branches within a specific time window, these statistics can depend on population structure that occurs outside the time-window of interest.

To address this limitation, we introduce a framework of **coalescent-based time-stratified statistics**, in which we explicitly consider *genealogical* events—as opposed to mutational events—that occur within a target time interval. By reformulating classical summary statistics in terms of haplotype-level allele sharing and expressing their expectations as functions of interval-specific coalescence probabilities, our approach correctly captures population structure at the intended temporal depth and avoids information leakage across time windows.

## 2 Results

### 2.1 Temporal covariance leakage in site-based and branch-based time-stratified statistics

We first use a simple example to illustrate a fundamental limitation of site-based and branch-based time-stratified statistics: even when they are defined using mutations or branches restricted to a specific time interval, they may fail to faithfully reflect population structure within that interval. Consider two populations that diverged at some time in the past, with no gene flow following the divergence. We focus on a time interval that is entirely more ancient than the divergence time, when the two populations were still part of a single ancestral population (Figure 2**A**).

**Figure 2:**
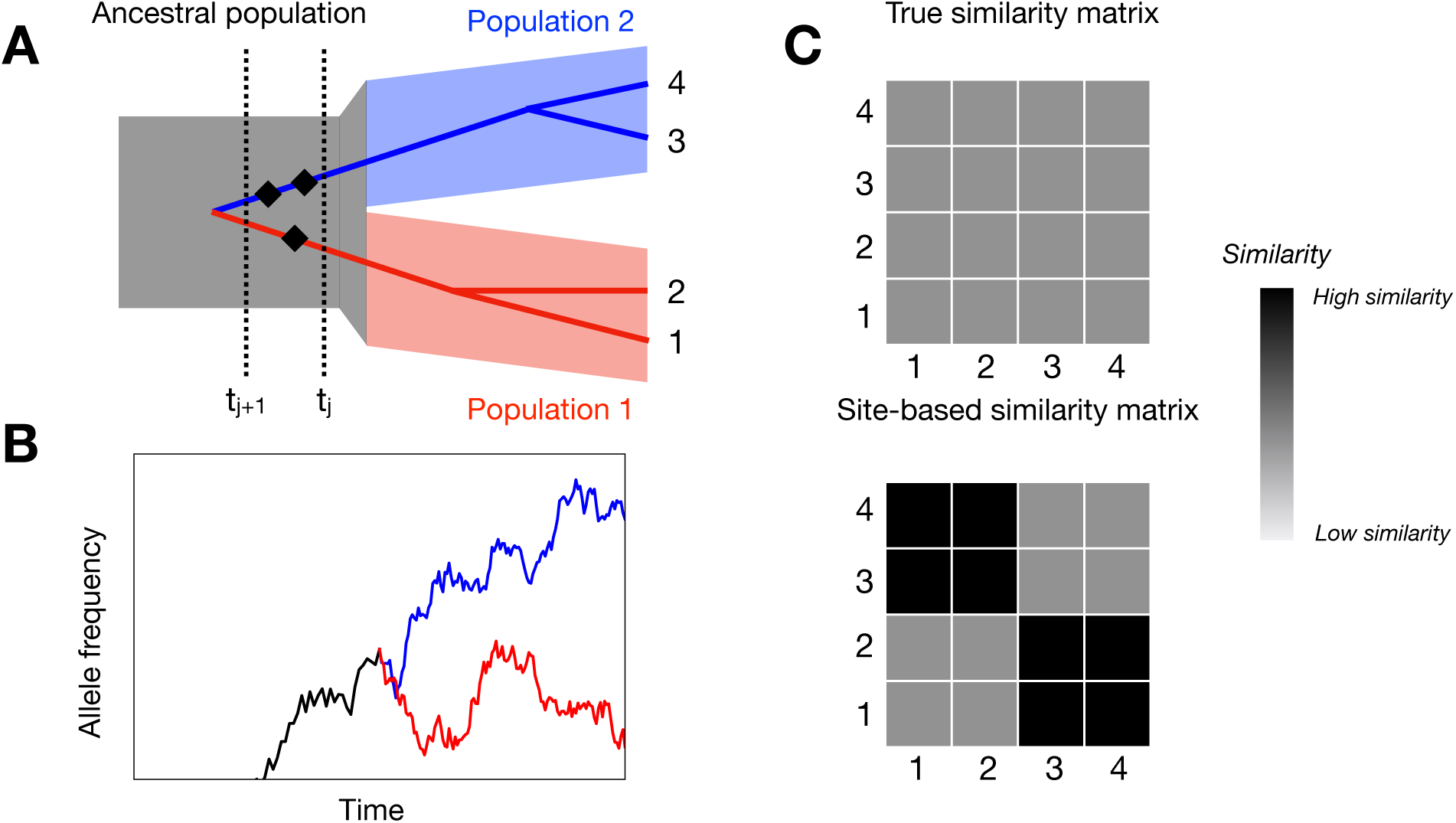
Temporal covariance leakage under a two-population clean-split model. (**A**) Backward-in-time view: sharing of mutations that arise within the focal time interval [*t_j_*, *t_j_*_+1_] can be driven by lineage coalescence occurring after the interval when the populations are isolated. (**B**) Forward-in-time view: allele frequencies of mutations arising in the ancestral population can diverge substantially after the population split due to genetic drift and lack of gene flow. (**C**) As a consequence, even when the focal time interval predates the population divergence and contains no true population structure, site-based time-stratified statistics may show higher similarity within populations than between populations.

From a **backward-in-time** perspective, samples from the same present-day population may share mutations that arose within the focal time interval (black diamonds in Figure 2**A**), not because those mutations represent population structure within the interval itself, but because the corresponding lineages have already coalesced in more recent time intervals after the population divergence. As illustrated in Figure 2**A**, lineages sampled from the same population are more likely to have coalesced before entering the ancestral population, whereas lineages from different populations are less likely to do so. Consequently, sample covariance induced by allele sharing at sites whose mutations arose in the ancestral interval can still be higher within populations than between populations, even though no population structure exists during the interval being examined (Figure 2**C**). In other words, covariance generated by more recent demographic history leaks into statistics intended to describe the ancestral interval.

The same phenomenon can be understood from a **forward-in-time** perspective. Even if a mutation arises in the ancestral population during the focal time interval, after the population split, its frequency will drift independently in the two populations due to genetic drift and the absence of gene flow. As a result, the same ancestral mutation may reach substantially different frequencies in samples drawn from the two populations (Figure 2**B**). Thus, although the mutation itself arose before any population structure existed, its present-day distribution and the induced covariance among samples (Figure 2**C**) reflect demographic processes occurring *after* the divergence.

This also makes clear that site-based statistics are throwing away information relevant to the time window. Even mutations arising before the intended time interval could be informative about population structure because of patterns of shared drift. In essence, the site-based definition reflects the present-day allele frequencies of mutations which arose in the past, but the allele frequency dynamics are affected by the demographic history and population structure from more recent times. In fact, several methods infer demographic history using only SNPs shared among populations, which likely arose well before any demographic events of interest [Pickrell and Pritchard, 2012, Patterson et al., 2012, Haak et al., 2015, Nielsen et al., 2023].

Although the discussion above is framed primarily in terms of allele-sharing in the site-based time-stratified statistics framework, the same reasoning applies directly to branch-based definitions. Branch-based statistics simply average over the placement of mutations along branches within a given time interval [Ralph et al., 2020]. As a result, they inherit the same problem and mainly differ in variance and computational efficiency.

Taken together, these two perspectives highlight the same underlying issue. Events occurring in more recent time periods can influence covariance in allele sharing patterns at sites whose mutations arose in older time intervals. As a result, site-based or branch-based time-stratified statistics computed for an ancient interval will also contain signals of more recent population structure. We refer to this phenomenon as **temporal covariance leakage**.

An important exception is when the time interval of interest begins at the present, i.e., [0, *t_u_*]. In this setting, there is no more recent time period whose effects can contaminate the statistic, and therefore site-based or branch-based time-stratified statistics restricted to these types of windows do not suffer from temporal covariance leakage and are suitable for studying recent population structure. This is exactly the approach taken in Twigstats [Speidel et al., 2025]. But, we again note that site-based versions of these methods fail to leverage the information present in older mutations.

### 2.2 Coalescent-based time-stratified statistic

From a backward-in-time perspective, the fundamental reason temporal covariance leakage arises is that allele sharing is induced by coalescence events that are more recent than the focal time interval. A natural way to eliminate this temporal covariance leakage is therefore to define time-stratified statistics **directly in terms of coalescence events occurring within the target interval**, rather than in terms of mutation sharing or branch sharing during that interval. We refer to statistics derived using this approach as **coalescent-based time-stratified statistics**.

Many summary statistics in population genetics can be written entirely in terms of second moments of individual or haplotype genotypes (e.g., the GRM) or allele frequencies (e.g., *F*-statistics). Throughout we focus on developing coalescent-based time-stratified versions of these second moments so that we can leverage these results for this class of statistics. For example, the (*i*, *j*)-th entry in the GRM can be written in terms of second moments as follows:

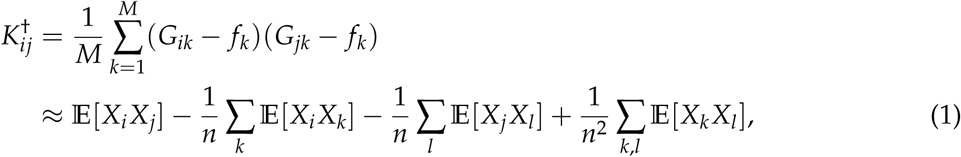

where *k* is the index for polymorphic sites among *M* total sites, *X_i_* denotes the allelic state at a randomly chosen genomic position in haplotype *i* among *n* total haplotypes, and expectations are taken with respect to the evolutionary process. As we will show below, we can decompose these second moments into contributions from coalescence events within each time interval.

Here we assume a bi-allelic mutation model, where the allelic state switches between 0 and 1 with mutation rate *µ*, and we also allow recurrent mutations. Although our derivations are presented under this model, they extend naturally to other mutation models because summary statistics including the GRM and *F*-statistics are invariant to allele polarization.

For two distinct haplotypes *i* ≠ *j*, the random variable *X_i_X_j_* equals 1 if and only if two conditions are satisfied: (1) *X_i_* = 1, which occurs with 50% probability under the bi-allelic model by symmetry; and (2) an even number of mutations occurs on the path connecting *i* and *j*. This condition is illustrated schematically in Figure 3.

**Figure 3:**
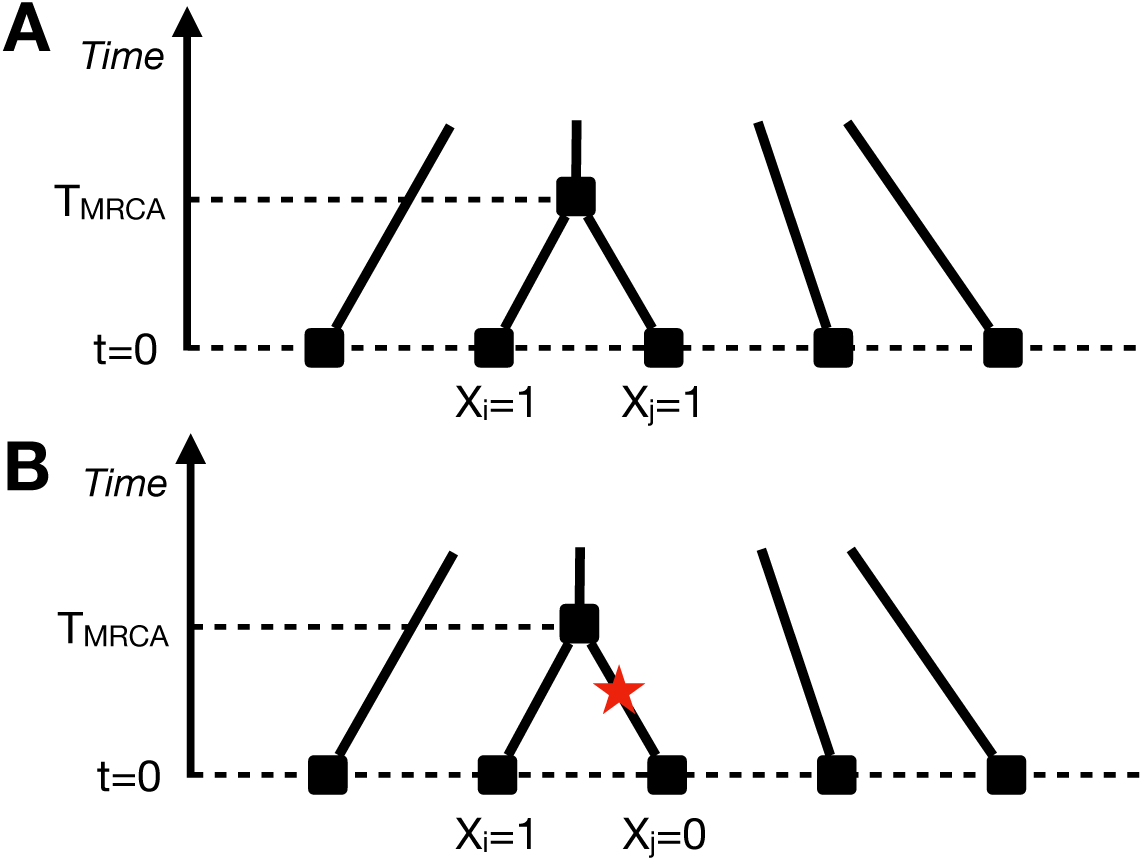
Conditions for *X_i_X_j_* = 1. (**A**) In order for *X_i_X_j_*to be 1, *X_i_* must be 1 and there must be an even number of mutations along the branches separating *i* and *j*. (**B**) If there are an odd number of mutations separating *i* and *j*, then *X_i_X_j_*must be 0.

As such, E[*X_i_X_j_*] can be computed as follows:

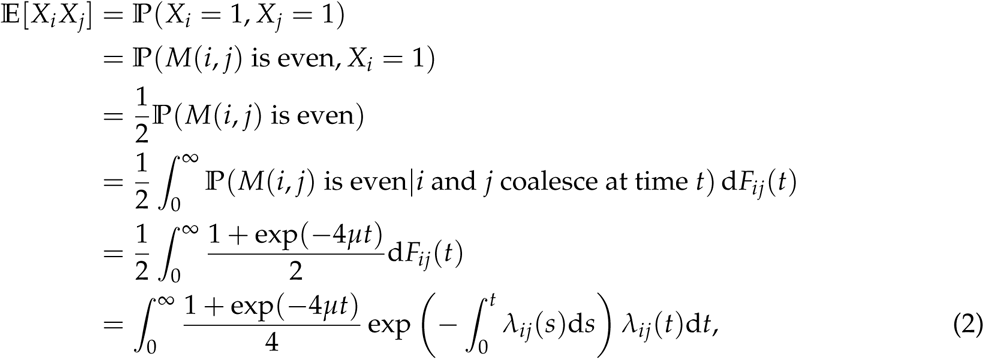

where *M*(*i*, *j*) is the number of mutations on the path from *i* to *j*, and *λ_ij_*(*t*) is the coalescence rate between lineage *i* and *j* at time *t*, and *F_ij_*(·) is the cumulative distribution function (CDF) of the pairwise coalescence time between *i* and *j*.

Equation 2 reveals that E[*X_i_X_j_*] can be stratified by the time at which a *coalescence* event occurs. In particular, the integrand in Equation 2 can be broken down into three parts. The first component, (1 + exp(−4*µt*))/4 does not depend on the demographic process in any way. The next component, exp(− ∫_0_*^t^ λ_ij_*(*s*)d*s*), reflects the probability that lineages do not coalesce with each other up until time *t*, which is entirely driven by what happens in the time interval [0, *t*). That is, this term purely reflects temporal covariance leakage. Finally, *λ_ij_*(*t*)d*t* can be thought of as the probability of coalescing in the infinitesimal interval *t* + d*t given that the lineages have not coalesced before time t*. This final term is the only term that reflects demographic forces at time *t* and by the Markovian nature of the coalescent does not depend on any demographic forces more recent than *t*. As such, it is reasonable to say that the conditional probability of coalescing within a time window is how the demography at time *t* contributes to E[*X_i_X_j_*] at time *t*: if we knew these conditional coalescence probabilities for every moment in time, we could calculate E[*X_i_X_j_*].

In practice, this stratification into infinitesimally small time windows raises challenges when connecting theory to data. As such, we now consider breaking down the integral in Equation 2 into contributions from larger time intervals. For convenience, we will assume a constant rate of coalescence within each time interval, writing *λ_k_* for the coalescence rate of the *k*-th interval, [*t_k_*, *t_k_*_+1_]. We will also use the notation Δ*_k_* to mean *t_k_*_+1_ − *t_k_* Here, we see that

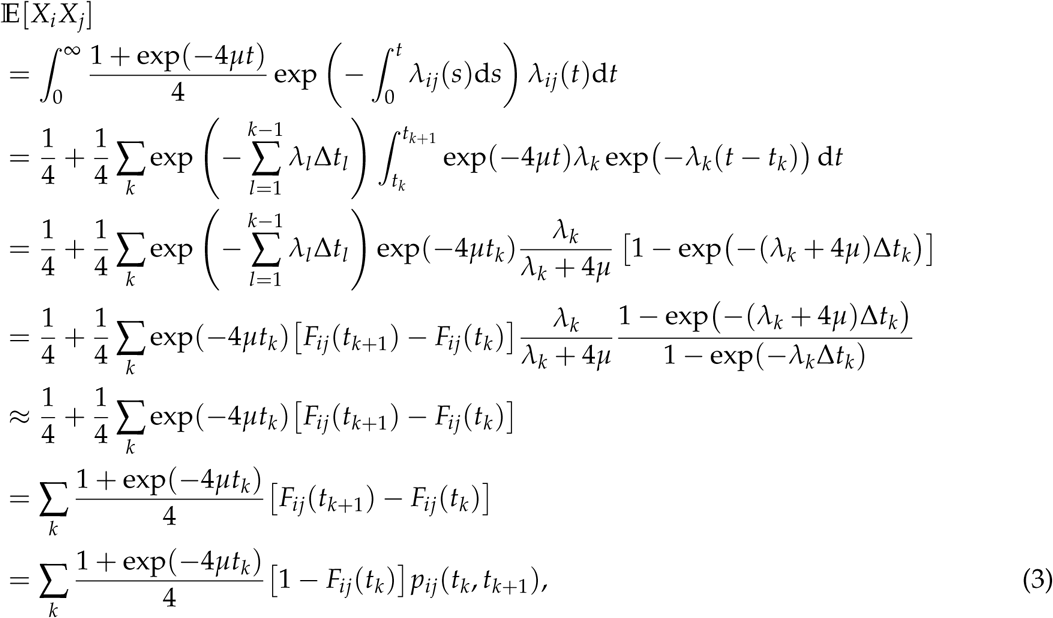

where the approximation assumes *µ* ≪ *λ_k_*, and *p_ij_*(*t_l_*, *t_u_*) is defined as the **conditional coalescence probability** within [*t_l_*, *t_u_*]:

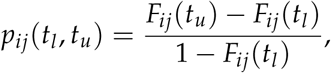

which represents the probability that lineages *i* and *j* coalesce within [*t_l_*, *t_u_*], conditioned on having remained un-coalesced until *t_l_*.

Again, we see that we can decompose E[*X_i_X_j_*] into contributions from coalescent events within different time windows, and within each time window there are three terms: 1) a mutational term independent of demography; 2) a temporal covariance leakage term that captures the probability of the lineages not coalescing until the start of the window; and 3) the contribution of coalescences within the window, conditioned on the lineages surviving to that point, which is the only term that depends on the demography within that time window, and it does not depend on the demography in any other time window.

We note that in the special case when *i* = *j*, *F_ii_*(*t*) = 1 for all *t*. This makes *p_ii_*(*t_l_*, *t_u_*) ill-defined, as would be expected as E[*X_i_X_i_*] = 1/2 under our model regardless of demography. As such, at no point in time does the demography contribute to these diagonal second moments.

These observations motivate us to define **coalescent-based time-stratified statistics** by retaining only the term that is informative about the population structure in the focal time interval, leading to the substitutions:

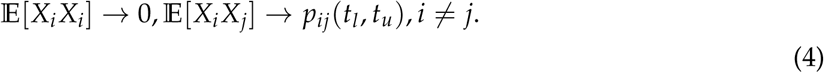

In practice, it is often useful to consider average coalescence probabilities across samples drawn from populations. For samples drawn from a population *A* of size *n*, we have:

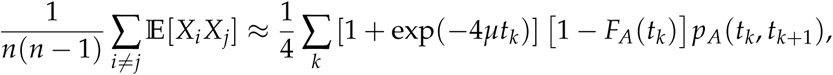

where the approximation requires the lineages in *A* to be exchangeable, *F_A_*(·) denotes the CDF of the coalescence time among a randomly chosen pair of lineages sampled from population *A*, and *p_A_*(*t_l_*, *t_u_*) represents the corresponding conditional coalescence probability. This can be conveniently computed using tskit through TreeSequence.pair coalescence counts() [Kelleher et al., 2018b, Ralph et al., 2020, Jeffery et al., 2026, N. Pope, personal communication]. Similarly, for samples drawn from two populations *A* and *B*:

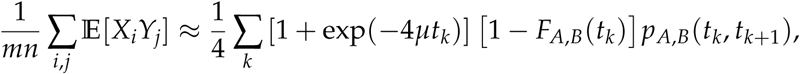

where again we assume that the lineages within *A* are exchangeable as are the lineages within *B*, and *F_A_*_,*B*_(·) and *p_A_*_,*B*_(*t_l_*, *t_u_*) are respectively the cumulative and conditional coalescence probabilities for a randomly chosen pair of lineages where one lineage comes from *A* and one from *B*. This can be also computed using tskit through TreeSequence.pair coalescence counts() [Kelleher et al., 2018b, Ralph et al., 2020, Jeffery et al., 2026, N. Pope, personal communication].

Similarly, this leads to the following substitutions when we have sample *X* from population *A* and sample *Y* from population *B*:

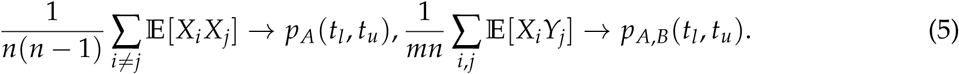

These substitutions provide a general recipe: when a summary statistic can be written in terms of second-order moments of haplotype genotypes, it can be reformulated as a function of interval-specific coalescence probabilities. In the following sections, we apply this framework to derive coalescent-based time-stratified versions of several commonly used population genetic statistics, and illustrate the utility of these statistics using ARGs simulated from realistic demographies.

### 2.3 Simulations

We used msprime [Kelleher et al., 2016, 2018a] to simulate ARGs across a range of evolutionary scenarios. All simulations were performed under two demographic models that capture distinct aspects of human population history.

The first model is the OutOfAfrica 3G09 model implemented in stdpopsim [Gutenkunst et al., 2009, Adrion et al., 2020]. This is a classical Out-of-Africa (OOA) model characterized by a sequence of population splits without subsequent admixture. It provides a simple, tree-like demographic history that serves as a baseline for evaluating time-stratified statistics in the absence of gene flow. For simplicity we remove the continuous migration between the populations, similar to the simulations in Fan et al. [2022].

The second model is an ancient Eurasian demographic model described by Pearson and Durbin [2023]. This model captures the complex population history of modern Europeans and their ancestral populations, incorporating both population splits and major admixture events. As such, it provides a contrasting setting to assess how different time-stratified statistics behave in the presence of gene flow and reticulate ancestry.

For convenience, we refer to these models as the *OOA model* and the *Eurasian model*, respectively, in the remainder of the paper. Throughout, we use demes to plot demographic models [Gower et al., 2022].

**Figure 4:**
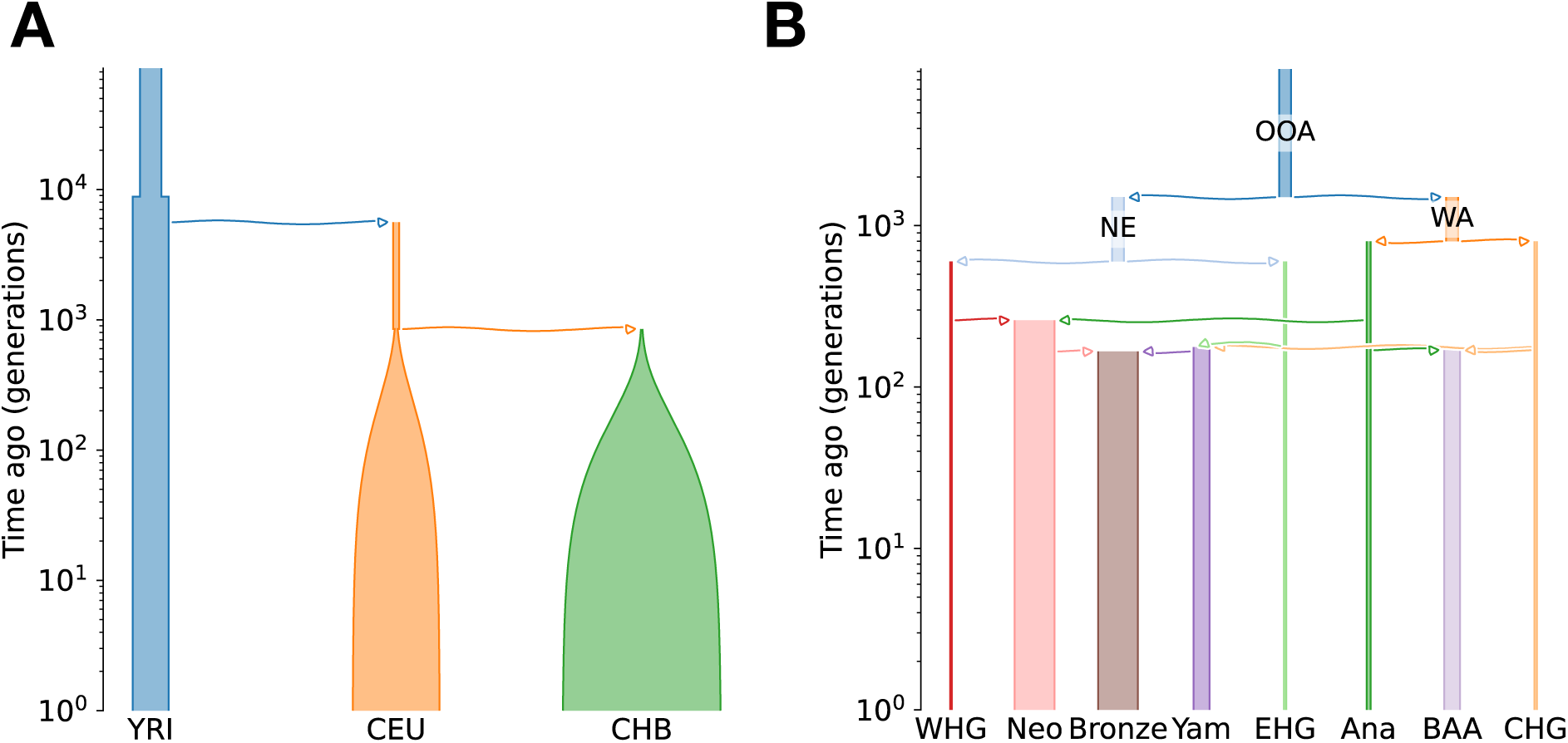
Demographic models used in this study. (A) The Out-of-Africa (OutOfAfrica 3G09) [Gutenkunst et al., 2009] model implemented in stdpopsim Adrion et al. [2020], featuring sequential population splits without admixture. (B) An ancient Eurasian demographic model from Pearson and Durbin [2023], incorporating both population splits and major admixture events.

### 2.4 Genetic Relatedness Matrix (GRM)

The GRM provides a natural first example for applying coalescent-based time-stratified statistics. For haplotype data, the centered but unscaled GRM is defined as:

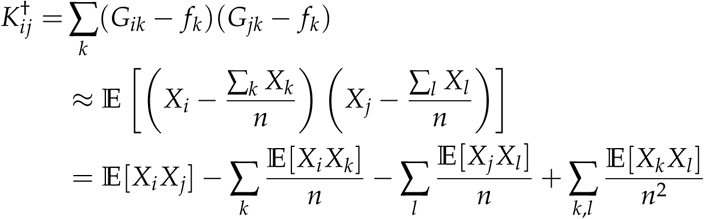

whose entries can be interpreted as the inner product between haplotypes *i* and *j*. By substituting the expectation terms according to (4), we have:

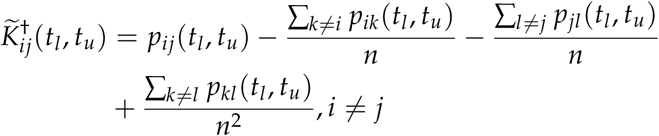

We intentionally use the GRM constructed using unscaled genotypes. Scaled genotypes are no longer second moments, but rather rational functions, falling outside the scope of our approach. Additionally, the idea behind using standardized genotypes is to up-weight rare variants and emphasize very recent structure, which is unnecessary when the goal is to isolate structure within specific time intervals.

To illustrate the behavior of the coalescent-based GRM, we used ARGs simulated under the OOA model to compute both the eGRM and the coalescent-based GRM in three time windows (Figure 5**A**): one postdating all splits ([0, 500] generations ago), one between splits ([1000, 1500] generations ago), and one predating all splits ([6000, 6500] generations ago).

**Figure 5:**
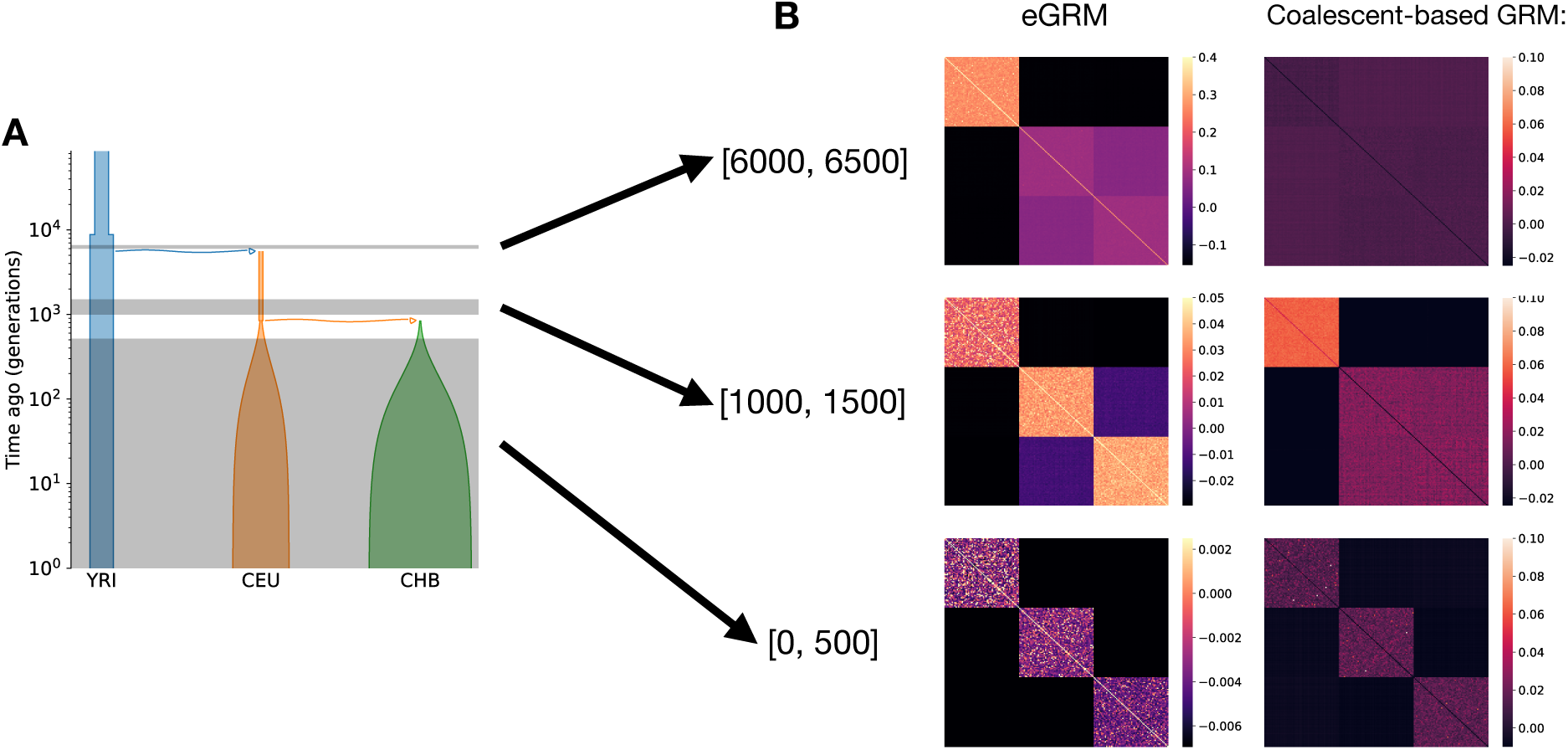
Comparison of eGRM and coalescent-based time-stratified GRMs. GRMs were inferred from the ARG under the OOA model. Shaded regions indicate the time intervals analyzed.

For time interval [0, 500], both the eGRM and the coalescent-based GRM capture the three-cluster structure (Figure 5**B**). However, for more ancient time intervals such as [1000, 1500], the eGRM assigns high covariance within CEU and CHB due to temporal covariance leakage, even though the time interval predates their split time (Figure 5**B**). Similar bias is also observed for the most ancient time interval, [6000, 6500], where the coalescent-based GRM captures the panmictic structure whereas the eGRM detects false structure induced by temporal covariance leakage (Figure 5**B**). Meantime, the coalescent-based GRM correctly represents the population structure in these three time intervals.

### 2.5 *F*-statistics

*F*-statistics are a widely used framework for studying population divergence and admixture. Unlike the GRM, which summarizes pairwise similarity among individuals, *F*-statistics are typically defined at the population level.

We begin with the simplest case, the *F*_2_ statistic, which measures allele-frequency divergence between two populations *A* and *B*:

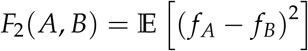

where *f_A_*and *f_B_* denote allele frequencies in the two populations. Although this definition is usually presented in terms of allele frequencies, it can be rewritten as a quadratic form in haplotype-level allele states. This reformulation allows us to substitute expectations involving haplotype states with coalescent probabilities, following (4):

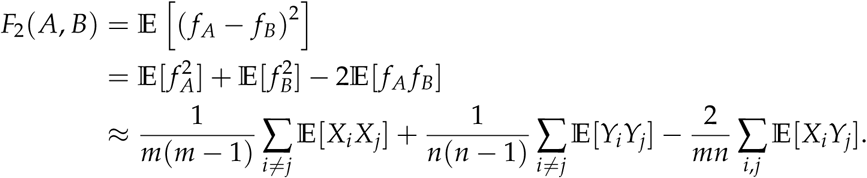

where the final line holds in the large-population limit, allowing us to neglect dependencies from sampling *m* lineages from population *A* without replacement, and likelwise for the *n* lineages from population *B*.

Substituting terms according to (4) yields the **coalescent-based time-stratified** *F*_2_ **statistic**:

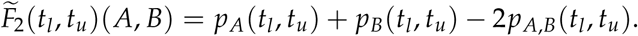

This expression admits a clear genealogical interpretation. When two populations are well mixed during the interval, within- and cross-population coalescence probabilities are similar, and 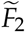 is close to zero. When populations are isolated, cross-population coalescence is suppressed relative to within-population coalescence, leading to a large positive value of 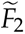.

It is important to note that *F*_2_(*A*, *B*) is designed to quantify the shared genetic drift between populations *A* and *B*, but our coalescent-based time-stratified 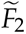(*A*, *B*) instead quantifies the shared drift between the *ancestors* of present-day populations *A* and *B*. Since only present-day genomes are sampled, these ancestral populations are represented by the ancestors of the sampled individuals. If there is gene flow or admixture, individuals presently in population *A* may have ancestors in both population *A* and *B* or even some other population. Our time-stratified statistic, 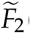(*A*, *B*), measures the amount of shared genetic drift that occurs within a time interval among those sets of ancestors, not necessarily among the populations themselves.

To compare the behavior of site-based and coalescent-based *F*_2_ statistics, we compute them using ARGs simulated under the OOA model for two pairs of populations: (YRI, CEU) and (CEU, CHB).

Site-based *F*_2_ remains nonzero even for intervals predating population divergence, reflecting post-divergence drift rather than ancestral structure (Figure 6**B**). In contrast, the coalescent-based 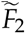 drops to zero once the time interval extends beyond the divergence time (Figure 6B).

**Figure 6:**
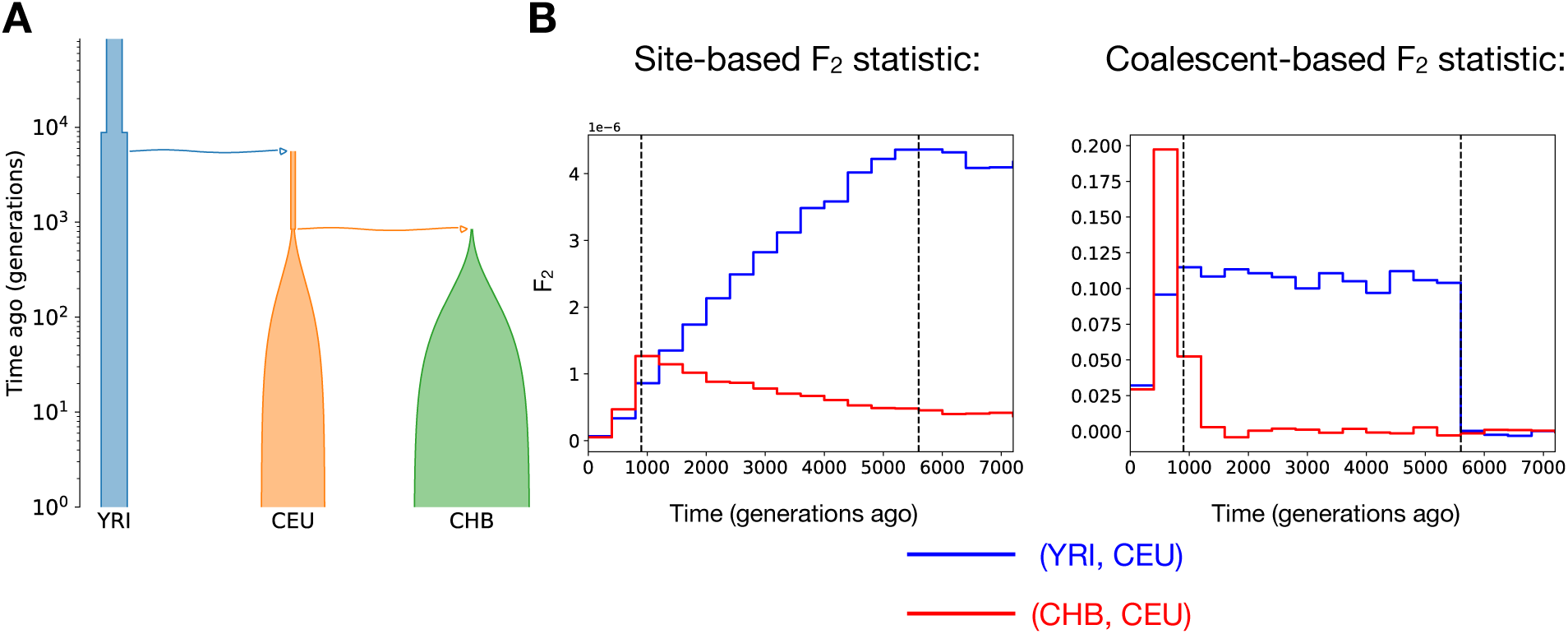
Comparison of site-based and coalescent-based *F*_2_ statistics. Statistics were inferred from the ARG under the OOA model for the population pairs YRI–CEU (blue) and CHB–CEU (red). Dashed vertical lines indicate the divergence times between CHB and CEU (left) and between YRI and CEU (right).

This can be explained by the temporal covariance leakage in the forward viewpoint (Figure 2**B**): even if a mutation arose in the past, the frequencies in two populations might still differ due to independent post-divergence drift.

### 2.6 Extension to *F*_3_ and *F*_4_

As *F*_3_ and *F*_4_ statistics can be viewed as linear combinations of *F*_2_ terms [Peter, 2016], the coalescent-based *F*_3_ and *F*_4_ follow immediately:

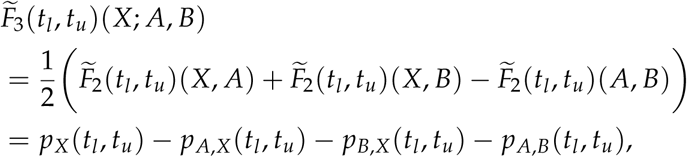

while the coalescent-based *F*_4_ statistic is

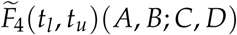

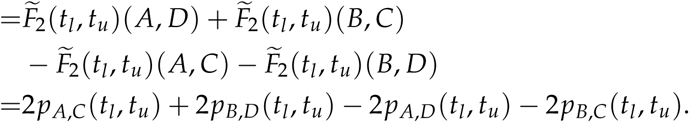

*F*_3_ statistics are less widely used than *F*_4_ statistics, so we focus on *F*_4_ for the remainder of this section. In practice, *D* is often chosen as a sufficiently diverged outgroup when computing *F*_4_(*A*, *B*; *C*, *D*). In these cases, *F*_4_ is used to test for admixture and can be interpreted as the extent of deviation from “treeness” in terms of the violation of the four-point condition from phylogenetics [Peter, 2016]. In this setting, coalescence between *D* and any ingroup lineage occurs far deeper in time than the time windows of interest. Over those windows, the relevant interval-specific coalescence probabilities involving *D* are therefore negligible, so the coalescent-based expression simplifies. In this common setting, the coalescent-based *F*_4_ essentially reduces to:

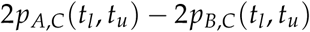

Recall that *F*_4_(*A*, *B*; *C*, *D*) can be understood as measuring the difference between mutations of *ABBA* and *BABA* patterns, and it has connections with coalescence probabilities. In the *ABBA* configuration, the lineages from *B* and *C* coalesce within the time interval before either joins *A*; in the *BABA* configuration, *A* and *C* coalesce first. Given either configuration, a mutation that occurs on the appropriate internal branch (i.e., the branch ancestral to the coalesced pair but not to the other ingroup lineage) produces an *ABBA* or *BABA* site pattern (Figure 7). Thus, the coalescent-based *F*_4_ corresponds exactly to the difference between the *BABA*-producing and *ABBA*-producing coalescent configurations.

**Figure 7:**
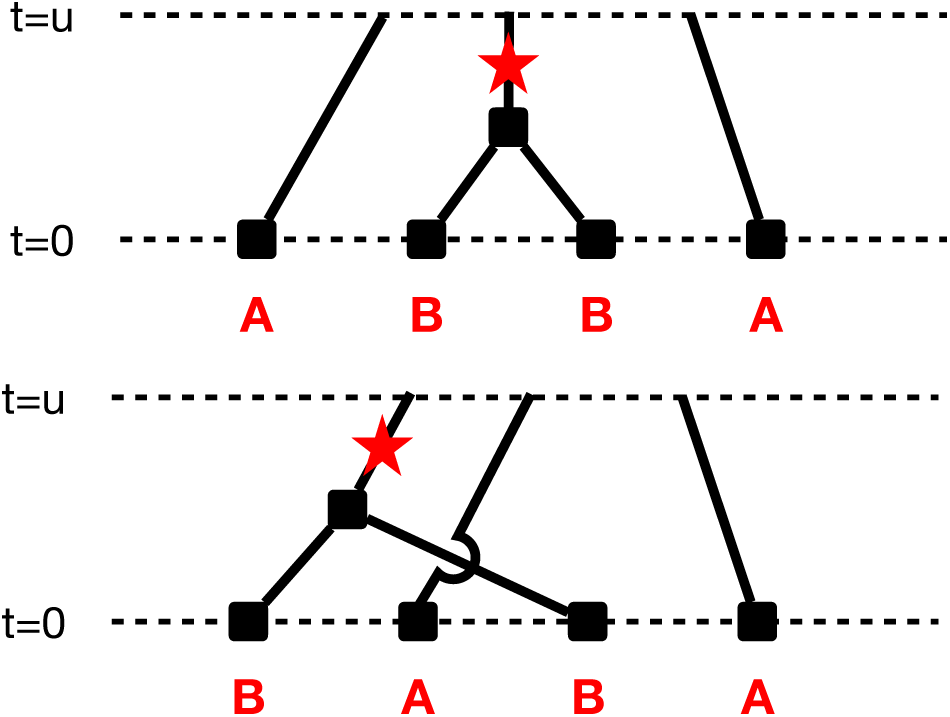
Coalescent configurations that give rise to *ABBA* and *BABA* mutation patterns. The four lineages correspond to the four populations used in the *F*_4_ statistics.

We next investigate how site-based and coalescent-based time-stratified *F*_4_ behave on simulated ARGs under the ancient Eurasia model. We analyze two quartets extracted from the full model: (EHG, WHG; Ana, CHG) and (Ana, BAA; CHG, EHG). The first quartet follows a purely tree-like history (no admixture among these four populations), whereas the second quartet includes an admixture event from CHG into BAA.

To understand the site-based behavior, recall that the classical definition,

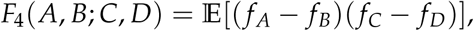

depends on how allele frequencies have drifted in each population after a mutation arises. For the tree-like quartet (EHG, WHG; Ana, CHG), the demographic model implies that the contrast (*f_EHG_* − *f_WHG_*) is generated only by drift occurring after EHG and WHG split, and similarly (*f_Ana_* − *f_CHG_*) is generated only after Ana and CHG split. Because these two splits lie on different branches of the tree and there is no gene flow connecting them, the two frequency contrasts are independent, and the site-based *F*_4_ remains near zero across time windows, as expected for a strictly tree-structured relationship (Figure 8).

**Figure 8:**
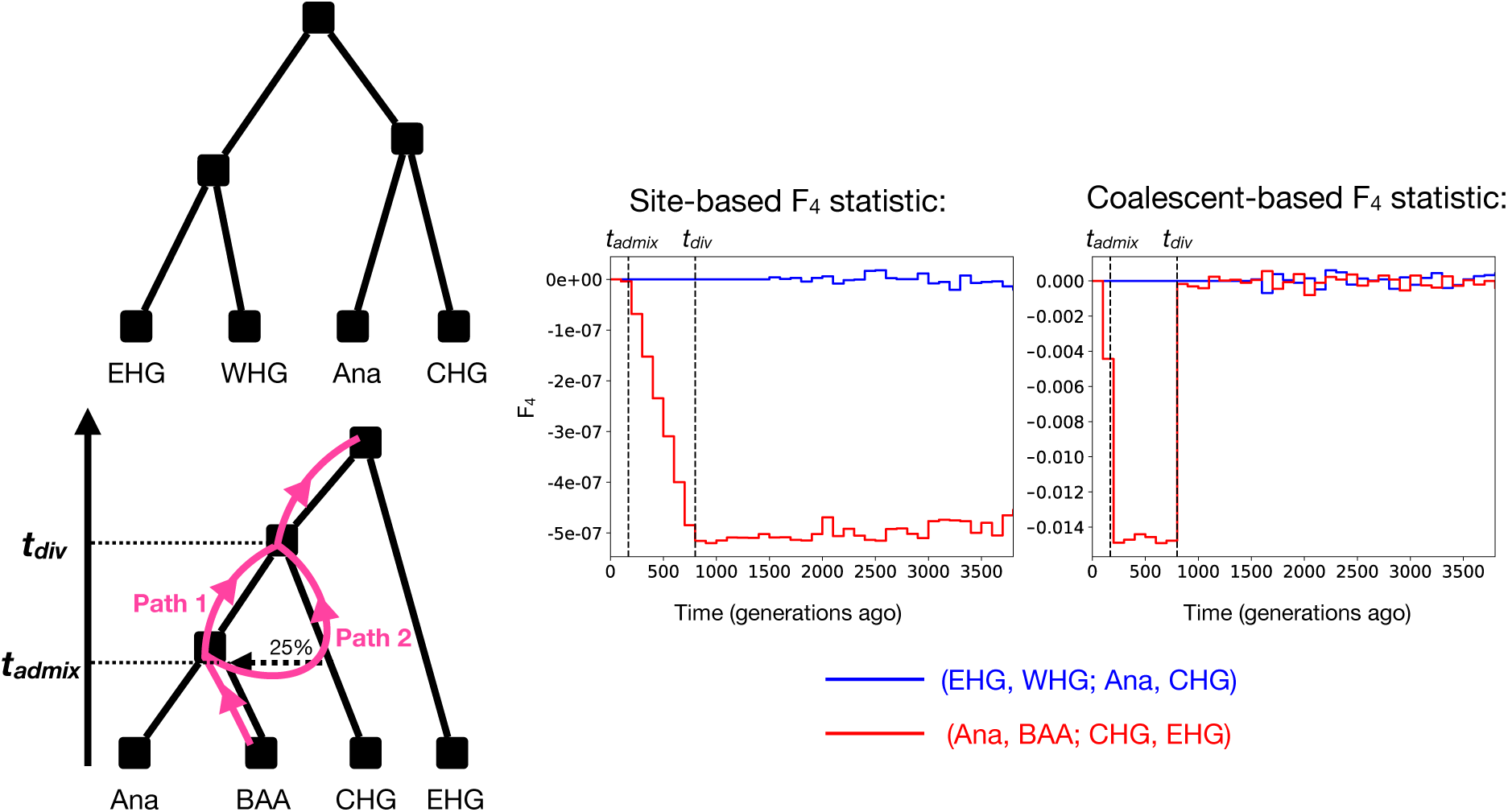
Comparison of site-based and coalescent-based *F*_4_ statistics. Statistics were inferred from the ARG under the Eurasian model for two population quartets: (EHG, WHG; Ana, CHG) and (Ana, BAA; CHG, EHG). Dashed vertical lines indicate the admixture time from CHG into BAA and the divergence time between Ana and CHG.

In contrast, for the admixed quartet (Ana, BAA; CHG, EHG), the two contrasts (*f_Ana_* − *f_BAA_*) and (*f_CHG_* − *f_EHG_*) are no longer independent for mutations that arose **before** the admixture event. The key observation is that BAA inherits ancestry from CHG, so—conditional on a mutation that predates admixture—allele frequencies in BAA and CHG are statistically coupled. The two contrasts are therefore negatively correlated. This explains the characteristic time pattern of the site-based *F*_4_: for windows containing only mutations that arose after the admixture, the four populations drift independently, and the statistic is close to zero; as we include older mutations that predate the admixture, the induced coupling between *f_BAA_* and *f_CHG_* drives *F*_4_ away from 0. While this site-based behavior is consistent with the demographic model, it is hard to interpret: the statistic in a given window reflects not only what happened within that interval, but also how allele frequencies subsequently drifted and mixed. The coalescent-based formulation makes the temporal interpretation more direct. With a proper outgroup *D*, the coalescent-based *F*_4_ can be read primarily as a comparison between *p_A_*_,*C*_(*t_l_*, *t_u_*) and *p_B_*_,*C*_(*t_l_*, *t_u_*), i.e., whether *C* is more likely to coalesce with *A* or with *B* within the interval.

For the (Ana, BAA; CHG, EHG) quartet, consider the time interval between (i) the admixture from CHG into BAA and (ii) the divergence between Ana and CHG. In this epoch, Ana and CHG are still distinct lineages in the model (so *p_Ana_*_,*CHG*_ = 0), whereas BAA carries CHG ancestry, so a lineage sampled from BAA has a non-negligible chance to coalesce with CHG within the interval (so *p_BAA_*_,*CHG*_ *>* 0). Consequently, the coalescent-based *F*_4_ is predictably negative within that temporal interval, and returns to approximately zero outside it.

We note that *F*_4_ statistics are often used to detect the deviation from “treeness” in the demographic history of the tested population quartet. However, the normal site-based *F*_4_ does not tell us any information regarding the temporal localization of the violation. In a tree-shaped demographic model, when tracing the ancestral lineages from one individual, there is only one unique path of historical populations that the lineages will go through. However, for admixed populations, because of the admixture event, there are 2 possible paths, and those paths differ only between the divergence time of the donor ancestries and the admixture event time (Figure 8). Equivalently, the ancestral lineages from the same individual can be distributed in two ancestral populations at the same time, which is not allowed in a tree-shaped demographic model. In other words, the violation to treeness only exists within this temporal interval, which our time-stratified coalescent-based *F*_4_ precisely captures and localizes.

### 2.7 *D*-statistics

Unnormalized statistics, such as *F*_4_, can be difficult to interpret quantitatively because their absolute values depend on factors unrelated to admixture, including heterozygosity and SNP ascertainment. This problem is even more severe when it comes to time-stratified statistics, as the length of the time interval can also affect their magnitude. As a result, while the sign of *F*_4_ is often informative, its magnitude is not directly comparable either across time intervals or across different studies.

To address this, it is common to use a normalized version, known as the *D*-statistic, which rescales *F*_4_ by its maximum possible value. This normalization yields a statistic bounded between −1 and 1, allowing values to be interpreted as the relative excess of allele sharing in one direction versus the other, and facilitating comparison across genomic regions and datasets.

Applying the same normalization principle to our coalescent-based formulation, we define the **coalescent-based, time-stratified** *D***-statistic** for a time interval [*t_l_*, *t_u_*] as:

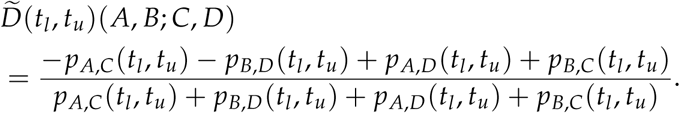

The temporal behavior of the coalescent-based *D*-statistic closely mirrors that of the coalescentbased *F*_4_, but with the important advantage that values are constrained to lie in [−1, 1] (Figure 9). This normalization makes it easier to compare the strength of signals across time windows and between different population quartets. In particular, intervals with no excess allele sharing yield values near zero, while intervals corresponding to admixture events show pronounced deviations whose sign reflects the direction of gene flow.

**Figure 9:**
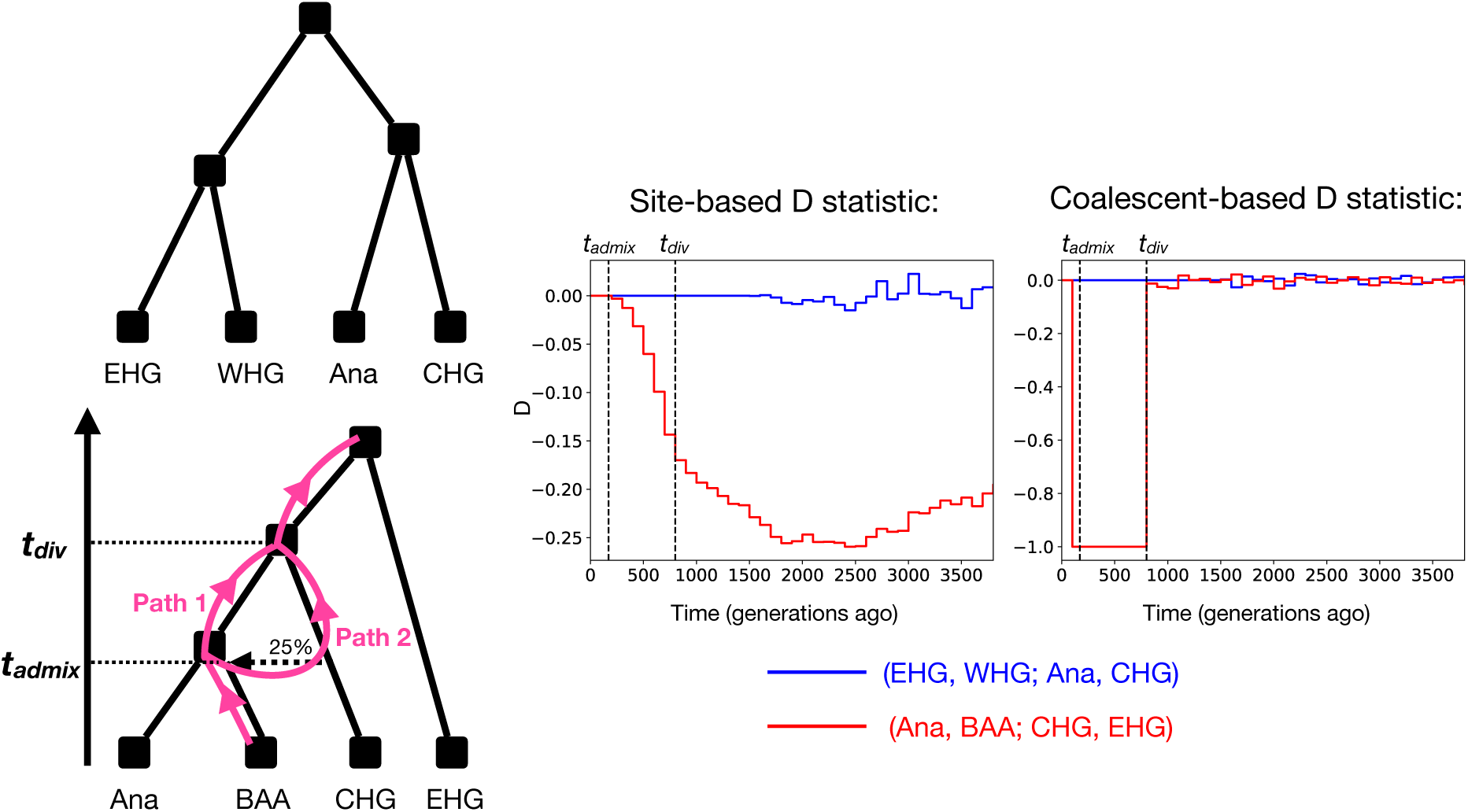
Comparison of site-based and coalescent-based *D*-statistics. Statistics were inferred from the ARG under the Eurasian model for two population quartets: (EHG, WHG; Ana, CHG) and (Ana, BAA; CHG, EHG). Dashed vertical lines indicate the admixture time from CHG into BAA and the divergence time between Ana and CHG.

### 2.8 Normalized *F*_2_ statistic

Similar to *F*_4_, *F*_2_ is also unnormalized, and its magnitude can be hard to interpret quantitatively, especially in a time-stratified setting. Here we introduce a normalized version of 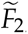, which we denote by 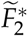, defined as:

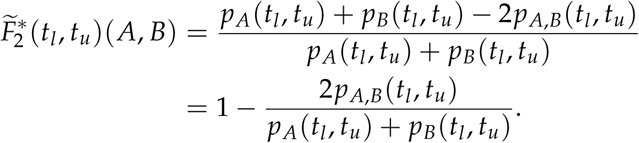

The normalized 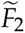 statistic 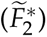 takes values between 0 and 1 and has a straightforward interpretation. It will be 1 when the two populations are completely isolated during [*t_l_*, *t_u_*], in which case lineages from different populations cannot coalesce and *p_A_*_,*B*_(*t_l_*, *t_u_*) = 0. Conversely, it equals 0 when populations *A* and *B* are panmictic during the interval, in which case within- and cross-population coalescence probabilities are equal, i.e., *p_A_*(*t_l_*, *t_u_*) = *p_B_*(*t_l_*, *t_u_*) = *p_A_*_,*B*_(*t_l_*, *t_u_*).

We next compute the time-stratified coalescent-based 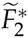 statistic under the OOA model, and compare it with another commonly-used site-based, normalized measure of population divergence, *F_st_*. For both population pairs considered—(YRI, CEU) and (CEU, CHB)—the site-based *F_st_* increases monotonically when tracing deeper back in time, without a clear correspondence to the population divergence time (Figure 10**B**). In contrast, the coalescent-based 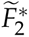 exhibits a much clearer temporal pattern: it is close to 1 for time intervals more recent than the population divergence and drops sharply toward zero for older intervals that predate the split (Figure 10**B**).

**Figure 10:**
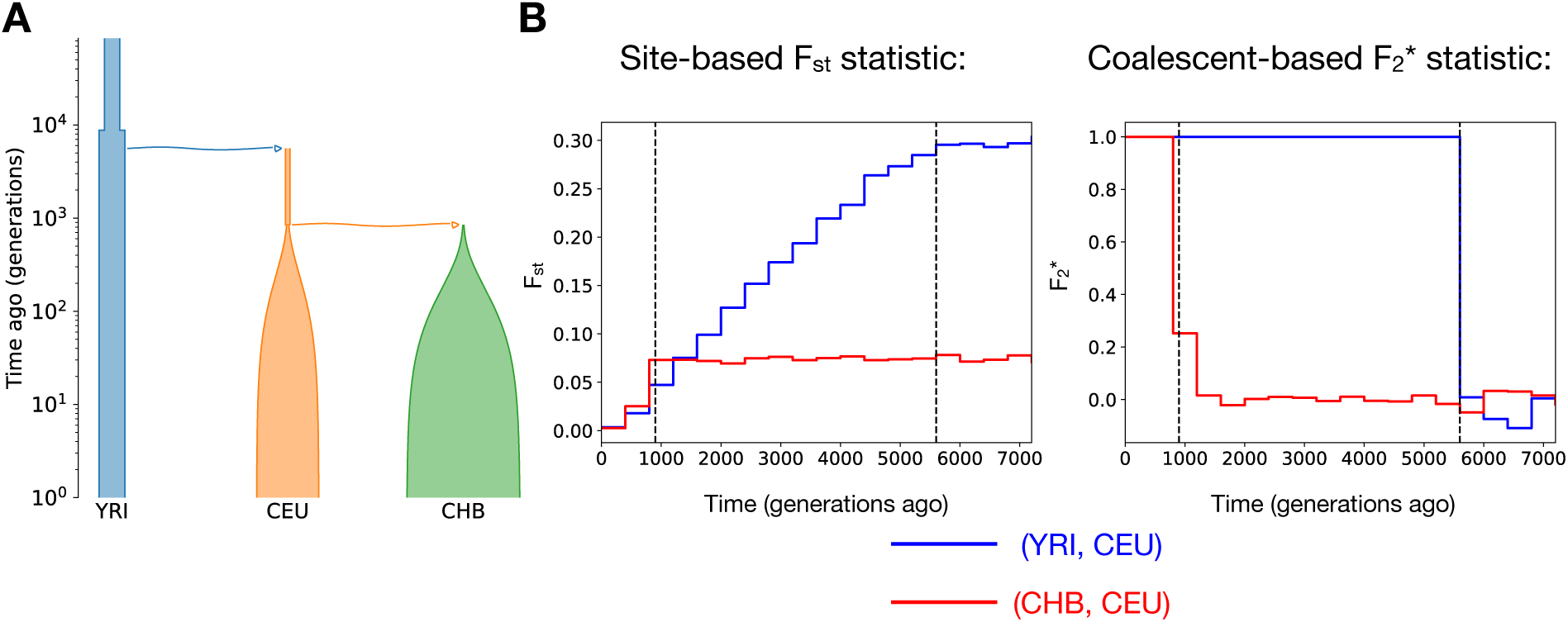
Comparison of site-based *F_st_* and coalescent-based 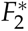 statistics. Statistics were inferred from the ARG under the OOA model for the population pairs YRI–CEU (blue) and CHB–CEU (red). Dashed vertical lines indicate the divergence times between CHB and CEU (left) and between YRI and CEU (right).

## 3 Discussion

In this work, we investigate the potential and challenge of defining time-stratified statistics from the ARG. We showed that time-stratified statistics derived by filtering mutations by age (site-based) or by restricting to branches in a time interval (branch-based), do not reliably isolate information from the intended time interval. Even when mutation ages are correctly restricted, both approaches can retain substantial contributions from more recent evolutionary history, leading to spurious signals of population structure in ancient time intervals. We refer to this effect as **temporal covariance leakage**.

This temporal covariance leakage arises because allele sharing among present-day samples reflects not only when mutations occurred, but also how lineages subsequently coalesced and, equivalently, how allele frequencies drifted after those mutations arose. As a result, time-stratified site- or branch-based statistics may capture demographic structure from more recent time periods, even when the analysis is nominally restricted to older epochs. This phenomenon highlights a fundamental limitation of time stratification strategies that condition only on mutation ages or branch locations, without explicitly conditioning on the underlying genealogical process.

To address this issue, we introduced **coalescent-based time-stratified statistics**, which reformulate classical summary statistics in terms of haplotype-level allele sharing and then express their expectations as functions of coalescence probabilities within a specified time interval. By conditioning on non-coalescence prior to the interval of interest, these statistics depend exclusively on genealogical events occurring within that interval, thereby fully removing contributions from more recent coalescence and allele-frequency drift. Using this framework, we derived coalescent-based definitions for the GRM, *F*-statistics, the *D*-statistic, and developed a new normalized version of 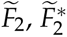. Each of these statistics reduces to simple combinations of interval-specific pairwise coalescence probabilities.

Through simulations under both population divergence and admixture models, we demonstrated that coalescent-based statistics recover population structure at the appropriate temporal depth, while site-based statistics can produce misleading signals that persist across time windows. These results collectively show that temporal interpretability in ARG-based analyses requires statistics that are explicitly defined in terms of coalescent events, rather than mutation ages alone. As large-scale genealogical inference becomes increasingly routine, coalescent-based time-stratified statistics provide a principled and flexible foundation for studying demographic history, population structure, and admixture across evolutionary timescales.

Although our coalescent-based statistics offer clear advantages for temporally localized demographic inference, there are applications for which site-based or branch-based statistics may be more appropriate. For example, in settings such as Genome-Wide Association Studies (GWAS) that aim to partition phenotypic variance across time, branch-based approaches like the eGRM may be more relevant. In particular, in that setting, one could imagine that variants of different ages may have different expected contributions to the phenotype. Thus, one really does care about all of the mutations arising within a time window as opposed to the demographic forces happening at that time. Whether coalescent-based, site-based, or branch-based statistics are best suited to a particular application therefore depends on the specific objective.

Our framework applies to second moments of either haploid allelic states or second moments of allele frequencies. Extending this coalescent-based approach to a broader class of summary statistics—including those involving higher-order moments or more complex dependencies— remains an important direction for future work.

We note that our framework reconstructs population structure only through the ancestors of present-day sampled populations. Consequently, historical populations that do not have any direct descendants among the sampled populations cannot be explicitly represented. Instead, their genetic contributions can only be inferred indirectly through the descendants that inherited ancestry from them. This limitation is particularly relevant for extinct or unsampled “ghost” populations that have shaped present-day genetic diversity, such as Neanderthals [Green et al., 2010], Denisovans [Reich et al., 2010], and ancient human groups including European hunter-gatherers and Ancient North Eurasians [Lazaridis et al., 2014]. Incorporating ancient DNA into our framework would provide a natural avenue for directly recovering these historical population structures and extending the temporal resolution of our approach.

Finally, our results are currently demonstrated using simulated ARGs. However, the inference of ARGs is not perfect: for example, even when the ground truth model has clean splits, inferred ARGs tend to infer gradual changes of relatedness. This is similar to observations of over-smoothing in methods such as PSMC and MSMC [Li and Durbin, 2011, Schiffels and Durbin, 2014]. As methods for inferring ARGs from real genomic data continue to improve, we expect that the power and applicability of coalescent-based time-stratified analyses in real data for addressing real-world population genetic problems will further increase.

## 4 Data Availability

The python scripts for implementing our coalescent-based time stratified statistics and an example Jupyter notebook are hosted in this Github repository: https://github.com/YunDeng98/Coalescent-based-time-stratified-statistics/tree/main.

## 5 Acknowledgments

We thank Hanbin Lee for valuable discussion on this project; Yulin Zhang, Alvina Adimoelja, and Tami Gjorgjieva for helpful feedback on the manuscript; Nathaniel Pope for guidance on using tskit functions; and Peter Ralph for insightful comments on this work.

